# Deactivation and mislocalization of a kinase protein induced by a single amino acid mutation on the proton transport catalytic aspartic acid

**DOI:** 10.1101/282822

**Authors:** Yang Zhang

## Abstract

Rhoptry protein 18 (ROP18) is a major determinant of strain-specific virulence in *Toxoplasma gondii*. The kinase activity of ROP18 is required for acute virulence, while the aspartate in the catalytic loop of ROP18 is considered essential for phosphoryl transfer. We showed that a single amino acid mutation at the catalytic aspartate residue (D409A mutation) significantly altered ROP18 kinase activity *in vitro*, and abolished ROP18-mediated ATF6β degradation. Furthermore, the investigated single amino acid mutation in ROP18 led to alternation of subcellular localization of ROP18 protein. Structural modeling analysis suggests that these phenotypes might be associated with D409A mutation induced conformation changes in ROP18.

Our findings demonstrate that a single amino acid mutation on the proton transport catalytic aspartic acid induced conformational alternations in ROP18 resulting in functional changes associated with ROP18 protein.

## Introduction

*Toxoplasma gondii* is an obligate intracellular parasite that causes significant morbidity and mortality particularly among the newborn and the immunocompromised[1]. This apicomplexan parasite engenders the common parasitic disease toxoplasmosis in almost all warm-blooded animals [2]. Serologic prevalence data reveal that *T. gondii* infections are highly prevalent in human population throughout the world, affecting up to half of the world’s population by some estimates [3-5].

There are predominantly three major lineages, namely types I, II and III, among *T. gondii* in North America and Europe[6]. Most infections in humans are caused by type II *T. gondii*[*7*]. As in the laboratory mouse model, *T. gondii* display vastly different levels of virulence among the three lineages. Type I parasites are mostly lethal with an infectious dose of a single parasite, whereas type II strains have an intermediate virulence that varies from 10^2^ to 10^5^ parasites, and type III parasites are considered avirulent[8].

Genetic crosses among lineages I, II, and III were used to map the genetic loci responsible for virulence. ROP5[9], ROP18[10], and ROP16[11] were identified as the major determinants of strain-specific virulence.

ROP 18 is a major determinant of virulence in type I *T. gondii*, contributing to the lethal phenotype in type I lineage as well as the lack of virulence in types II and III parasites [12]. ROP18 localizes to the rhoptry bulb and it is a member of the ROP2 superfamily [13]. When parasites invade the host cells, rhoptries discharge ROP18 together with other ROP2 superfamily kinases to form nascent parasitophorous vacuoles[14] and some into host cell cytosol [15]. Additional studies demonstrate that the N-terminal arginine-rich domain of ROP 18 is critical to its function as a virulence determinant[16].

Kinase activity of ROP18 is required for acute virulence by inducing host protein phosphorylation and degradation. The aspartate in the catalytic loop (“HRD” motif) is considered as a catalytically essential residue responsible for phosphoryl transfer[17]. ROPs lacking one or more of those conserved catalytic motifs, are predicted to be catalytically inactive and have been termed pseudokinases[9]. ROP2/8, ROP4/7, ROP5, ROP22, ROP36, ROP40, and ROP42/43/44 are identified or likely pseudokinases[18,19]. Interestingly, the catalytic Asp in kinase-conserved “HRD” motif is replaced with a basic residue specifically in ROP4/7 (HGK), ROP5 (HG[R/K/H]), ROP22 (HTH), ROP36 (HGH), ROP40 (LRR) and ROP42–43-44 (HGK) [9,17]. Among them, ROP5’s mechanism of action is well understood; it adopts the kinase fold by binding ATP in a non-canonical conformation. Sequence alignment reveals strong similarity between ROP5 and ROP18, sharing about 28% of sequence identity[20].

ATF6β was identified as a main host target of ROP18; ROP18 phosphorylates ATF6β, thereby targeting it for proteasome-dependent degradation [21]. ATF6β is synthesized as an endoplasmic reticulum (ER)-transmembrane protein, which is activated during ER stress response (ERSR) [22]. ER stress-induced proteolysis of membrane-bound ATF6 liberates the N-terminal fragment of ATF6, which then translocates into the nucleus and activates its target ERSR genes[23].

It is postulated herein that mutating the catalytic aspartate residue of ROP18 might alter its conformation and function. Therefore, it is tested that a single amino acid mutation at the catalytic aspartate residue could significantly alter ROP18 kinase activity *in vitro*. Previous studies have demonstrated that this mutation could affect intracellular parasite development and virulence[24].

ATF6β protein level was abrogated in wild-type cells co-transfected with ROP18 but not in D409A mutants, demonstrating that kinase activity is involved in ROP18-mediated degradation. ATF6β degradation was precluded with the addition of protease inhibitors, indicating that this particular ROP18-mediated degradation is proteasome-dependent. Colocalization of ROP18 kinase dead and ATF6β suggested that their physical interaction occurs in the Golgi apparatus. Association of ROP18 with the cell membrane was observed, suggesting a membrane-targeting strategy for ROP18. Intriguingly, wild-type ROP18 remained membrane-bound, whereas ROP18 kinase dead was associated with intracellular trafficking vesicles and some ultimately secreted out of the cell. It is likely that changes in protein hydrophobicity or kinase activity contributed to ROP18 kinase dead’s aberrant secretory pattern. It is also possible that membrane attachment is imperative for *T. gondii’s* acute virulence. Collectively, these findings indicated that a single amino acid mutation on the proton transport catalytic aspartic acid induced functional changes in ROP18, which might be associated with conformation changes within the ATP binding pocket.

## Materials and Methods

### Bacteria Strains, Cell lines and Plasmids

BL21 (DE3) V2RpACYC-lic-LamP-phosphatase cells (a kind gift from Dr. Raymond Hui, Structural Genomics Consortium), E. cloni^®^ 10G cells (Lucigen, USA), XL1-Blue supercompetent cells (Agilent, USA).

Human Embryonic Kidney 293T cell (HEK 293T) was a kind gift from Dr. Paul Dear, MRC Laboratory of Molecular Biology, Cambridge, UK.

pcDNA ATF6β-YFP; pcDNA HA_ATF6β_YFP-T2ACFP(with a self-cleavage signal [T2A] in between to produce the same level of both proteins); pcDNA HA-tagged ATF6β expression vectors were obtained from Dr. Masahiro Yamamoto, Osaka University, Japan. The pGEX-6p-ROP18mat-His plasmid was a kind gift from Dr. David Sibley, Washington University. The pcDNA5/FRT/TO was a kind gift from Dr. Paul Dear, MRC Laboratory of Molecular Biology, Cambridge, UK. pCherry-C2 Rab11a and pCherry-LAMP1 mammalian expression vectors were obtained from Dr. George S. W. Tsao, Department of Anatomy, University of Hong Kong.

### Chemical Bacterial Transformation

50 μl of pre-thawed competent cells were incubated with 1 μl of plasmid on ice for 0.5 hours. Then the cells were heat shocked at 42 °C for 60 seconds and returned to ice for 2 minutes. 250μl room temperature SOC medium (0.5% yeast extract, 2% tryptone, 10 mM NaCl, 2.5 mM KCl, 20 mM MgSO_4_, 20 mM Glucose) was added to the heat-shocked transformed competent cells and incubated at 37 °C for 1 hr with shaking at 220 rpm. After incubation, 20 μl of transformation mix was spread on the LB agar plate (1% Tryptone, 0.5% Yeast Extract, 1%NaCl, 1.5% agar) containing the appropriate antibiotic. The plate was placed in the 37°C incubator and grown overnight.

### Recombinant Protein Expression and Purification

BL21 (DE3) V2RpACYC-lic-LamP-phosphatase bacteria cells were transformed by chemical transformation with different plasmids. Transformed bacterial clones were subsequently grown on 5ml of liquid LB (lysogeny broth) in the presence of 100μg/ml ampicillin and 30μg/ml chloramphenicol overnight. The bacteria culture was diluted 1:100 into 500ml Terrific Broth medium (1.2% Tryptone, 2.4% Yeast Extract, 0.94% K_2_HPO_4_, 0.22% KH_2_PO_4_, 0.5% Glycerol) containing 100μg/ml ampicillin and 30μg/ml chloramphenicol and grown on a rotator shaker at 200 rpm and 37°C to an OD600nm of 0.6. The protein expression was induced by addition of isopropyl-β-D-1-thiogalactopyranoside (IPTG; Sigma Aldrich) to a final concentration of 1 mM and the cell were grown on a rotator shaker at 180rpm at 18°C overnight. Cells containing the recombinant proteins were harvested by centrifugation at 5000g and 4°C for 10 minutes.

The cells pellets were weighed and resuspended, lysed with 2XCelLytic B lysis buffer (Sigma, USA) containing 1xcOmplete, Mini, EDTA-free Protease Inhibitor Cocktail Tablets (Roche), Benzonase (50 units/ml; Sigma, USA), lysozyme (0.1 mg/ml; Sigma, USA). The CelLytic B 2X to cell mass ratio should be 5 ml per gram of wet cell paste. Then, the lysate was centrifuged at 16000 g for 10min at 4°C.

Glutathione Sepharose 4B(GE Healthcare, Sweden) was washed by PBS (140mM NaCl, 2.7mM KCl, 10mM Na_2_HPO_4_, 1.8mM KH_2_PO_4_, pH 7.3) twice. The cell lysate was incubated with the prepared Glutathione Sepharose 4B for 2h at 4 °C using gentle agitation. The supernatant was carefully removed by centrifugation at 500g for 5 min and the beads were then washed with PBS three times. The bound GST protein was then eluted with elution buffer (50mM Tris-HCl, 10mMreduced glutathione, pH 8.0). Pierce BCA protein assay kit (Thermo Scientific, USA) was used to measure the protein concentration, which is based on bicinchoninic acid (BCA) for the colorimetric detection and quantitation of total protein.

### ADP-Glo™ Kinase Assay

ADP-Glo™ Kinase Assay is a luminescent ADP detection assay to measure kinase activity by quantifying the amount of ADP produced during a kinase reaction. The kinase activity of ROP 18 (WT) or ROP 18(D409A) were tested according to the manufacturer’s instructions.

### Protein Separation by Sodium dodecyl sulfate-polyacrylamide gel electrophoresis (SDS-PAGE) and Western blot

Samples for SDS-PAGE were made with sample loading buffer and sample reducing agent, and heated at 95°C for 6 min before loading onto a 4-15% pre-cast Mini-PROTEAN^®^ TGX™ Precast Gel (Bio-rad, USA), which was run at 15 V/cm for 60min. The running buffer was 25 mM Tris, 192 mM Glycine, 0.1 % SDS, pH 8.3.

After electrophoresis, the polyacrylamide gel was soaked in transfer buffer (25 mM Tris, 192 mM Glycine, 20 % methanol (v/v)). PVDF membrane (Hybond-P; Amersham Pharmacia Biotech, UK) was cut to the correct size for the gel and prepared for transfer by soaking in methanol. Sponges and blotting paper were soaked in transfer buffer. Place successively from the anode to the cathode: paper, the PVDF membrane, the gel, and further paper taking care to remove any bubbles between the different layers. The transfer cassette was loaded into the gel tank and connected to power supply for 3hr at 35V. Following transfer, the membrane was incubated with 5% skimmed milk prepared in TBST (TBS containing 0.1% Tween-20) for 1hr. After blocking, the membrane was incubated with appropriately diluted primary antibodies for 1hr at room temperature. Then the membrane was washed three times with TBST for 10min. The membrane was then incubated with the secondary antibody (against the primary antibody) for 1hr at RT. Then the membrane was washed three times with TBST and applied with mixed detection solution A and Bin the ratio of 40:1 from ECL Plus Western Blotting Detection Reagents (GE Healthcare, USA) for 1 min at room temperature. Excess reagent was removed and the x-ray film was developed, fixed and washed in the dark room. The protein size was estimated using Rainbow Molecular Weight Marker (GE Healthcare, USA).

### Coomassie Blue Staining

After electrophoresis, the polyacrylamide gel was released and fixed in fixing solution (50% methanol and 10% glacial acetic acid) for 1hr with gentle agitation. The gel was then stained in 0.1% Coomassie Brilliant Blue R-250, 50% methanol and 10% glacial acetic acid for 5hr with gentle agitation. The gel was destained in destaining solution (40% methanol and 10% glacial acetic acid) until the background is sufficiently reduced.

### ROP18 Mammalian Expression Vector Construction

Restriction digests of pGEX-6p-ROP 18 plasmid DNA were completed by using BamHI-HF™ and NotI-HF™ (NEB), according to the product manuals. The ROP 18 DNA fragment was excised and ligated with the pcDNA5/FRT/TO by using T4 DNA Ligase (NEB), according to the product manuals. Then the reaction mixture was transformed into E. cloni^®^ 10G competent cells. Individual colonies were picked and the plasmid was extracted for screening the incorporation of the pcDNA5/FRT/TO: ROP 18 DNA construct by restriction digestion with BamHI and NotI. The appropriate plasmids were sent for DNA sequencing to confirm the integrity of the plasmids. The sequencing primer is: 5’-GACTTGCAGAGGGAGTCGTC-3’.

### Site-Directed Mutagenesis

ROP18 D409A and ROP18 R223E mutant was generated by using QuikChange^®^

#### Site-Directed Mutagenesis Kit

The PCR amplification was performed according to manufacturer’s instructions.

The PCR product was treated with 10 units of Dpn I enzyme for 1 hour at 37°C to digest the parental nonmutated ROP18 plasmid. 1 μl of the Dpn I-treated plasmid was transformed into XL1-Blue supercompetent cells by heat shock then the transformation reaction were plated on a LB–ampicillin agar plate and incubated overnight at 37°C.

The next day, several colonies were picked and inoculated into 10 ml LB+ ampicillin medium, and incubated overnight at 37°C. The ROP18 D409A mutant plasmid was extracted by using the QIAprep Spin Miniprep Kit. The plasmids were sent for DNA sequencing to confirm the integrity of the plasmids by using the upstream primer.

### Cell Culture

Human Embryonic Kidney 293T cell line was grown in Dulbecco’s modified Eagle’s medium (DMEM, (Sigma Aldrich, USA)) supplemented with 10% fetal bovine serum (FBS, (Sigma Aldrich, USA)), 100U/ml penicillin-streptomycin (Sigma Aldrich, USA), and 1% L-Glutamine (Lifetech, USA). For cell maintenance, cells were first washed with Ca^2+^, Mg^2+^-free PBS and then detached by treatment with trypsin/EDTA (0.05 % trypsin, 0.2 % EDTA) at 37°C for several minutes. As soon as the cells detached from the cell culture flask, fresh medium containing 10% FBS was used to inactivate the trypsin and the detached cells were centrifuged at 500g for 5min. The pellet was resuspended in fresh medium containing 10% FBS and the above-mentioned antibiotics. The adequate dilutions of cells were seeded in new cell culture flask.

Mycoplasma contamination was monitored from time to time by using the MycoAlert™ Mycoplasma Detection Kit, which measure the luciferase signal from ATP formed by mycoplasmal enzymes.

### Mammalian Cells Transfection

For plasmid transfection, low passage of 293T cells (5×10^5^ cells/well) were seeded in 6-well plates one day before transfection in 10% FBS DMEM medium with antibiotics. On the next day, 2 μg of mammalian expression vectors were diluted in 100 μl of pure DMEM. After mixed and 5 min incubation at room temperature, 6 μl of Fugen HD transfection reagent (Promega) was added and the mixture was incubated for additional 15 min. Then the mixture was added on top of the cells drop by drop. After 24-48 hrs incubation at 37°C, the transfection efficiency was checked either by immunofluoresence or by Western blot analysis.

### Immunofluorescence Staining

293T cells grown on the glass cover slips were subjected to transfection treatment. At 24-48 h post-transfection, the cells were fixed with 4% paraformaldehyde (PFA) in PBS (15 min at RT). The fixed cells were washed with PBS 3 times and permeabilized with 0.3% Triton X −100 in PBS (15 min at RT). After washing with PBS, the permeabilized cells were incubated with PBS.BSA for 1 h at RT. Then the cells were incubated with primary antibodies against rabbit anti-ROP 18(1: 500) and mouse anti-Giantin (1:100) for 1 h. After 3 washes in PBS for 10 min each, cells were then incubated to goat anti-mouse Alexa Fluor^®^ 568 or goat anti-rabbit Dylight 350 conjugated immunoglobulin G (IgG) for 1 h. The cells were washed three times in PBS and the coverslips were mounted on the glass slide with mounting solution. For mitochondrial labeling, MitoTracker^®^ dyes were diluted directly in the growth medium at the concentration of 100nM and incubated for 30min at 37 °C. After incubation, the cells were fixed in ice-cold methanol for 15 min at −20°C then rinsed 3 times with PBS for 10 min each.

### Fluorescence reporter assay

The ATF6β-YFP/CFP reporter plasmids were transiently co-transfected with ROP18 (WT) or ROP18 (D409A) into 293T cells using Fugen HD transfection reagent (Roche). Fluorescence signal of transfected cells were measured using the microplate reader (BMG labtech). The fluorescence intensity levels of YFP were measured at an excitation wavelength of 500±10 nm and an emission wavelength of 530±10 nm, CFP were measured at an excitation wavelength of 430±10 nm and an emission wavelength of 480±10 nm.

### Immunoprecipitation

293T cells grown in 6 well plates were transfected with ATF6β together with ROP 18(WT) or ROP 18(D409A) plasmids for 48 h.

Cells were washed and resuspended in 0.5 ml of RIPA lysis buffer and protease inhibitor cocktail tablets (Roche). After centrifugation, the supernatant was incubated with anti-HA antibodies for 1 h. The immune complexes were recovered by adsorption to Dynabeads^®^ Protein A 1 h at room temperature. After five washes in lysis buffer, the immunoprecipitates were analyzed by western blot analysis.

### Protein secretion analysis

The cell culture medium was collected and centrifuged at 13000xg for 5 min to remove the cell debris. The medium was then concentrated by 30 kDa cutoff Amicon concentrators, followed the manufacturer’s instructions. The concentrated solutions were analysed by western blotimmunoblotting with anti-β-Tubulin and anti-ROP18 antibodies respectively. Non-transfection and vector only were used as the controls. The cell lysates from adherent culture cells were collected as transfection controls.

## Results

### Structural modeling of wild-type ROP18, ROP18 kinase dead, and homologies to known ROP18 crystal structure

The crystal structure of ROP18 kinase domain has been solved [25]. By using PDBeFold (http://www.ebi.ac.uk/msd-srv/ssm/), a web service for identifying similarities in 3D protein structures, a comparison between predicted structures of wild-type ROP18 or ROP18 kinase dead (D409A mutant) generated from I-TASSER and the known, existing 3D structure determined by x-ray crystallography (PDB code 4JRN) was made.

The alignment results between predicted structure and x-ray 3D structure of kinase domain are hereby listed (Table 1). The structural relatedness of the proteins involves consideration of average root-mean-square deviation (RMSD), Q score (CaG alignment) and % Sequence Identity.

The degree of superimposability could be extrapolated from RMSD. A low RMSD between predicted structure of wild-type ROP18 and the known x-ray structure would indicate a high degree of superimposability. However, a high RMSD was obtained when comparing predicted structure of ROP18 kinase dead to predicted wild-type ROP18 structure or the known x-ray structure, suggesting a big structural divergence from both predicted wild-type ROP18 and known ROP18 crystal structure.

Q-scores take into account the number of residues in matched secondary structure elements (SSEs) as well as positions in space. High Q-scores are obtained when comparing x-ray structure of predicted wild-type ROP18 with that of known ROP18 crystal structure. This observation suggests that a large number of residues in equivalent structural elements superimpose well in three-dimensional space. However, low Q-scores are observed when comparing predicted structures of ROP18 kinase dead to that of wild-type ROP18 or known ROP18 x-ray crystallography.

There is a 61% sequence homology between the predicted kinase dead structure and the x-ray 3D structure, where an 82% sequence identity between predicted wild-type ROP18 models and the existing x-ray 3D structure.

The high degree of similarity between the wild-type ROP18 prediction, along with the known x-ray 3D structure, indicated that I-TASSER prediction results were reliable. Furthermore, D409A mutation could cause significant structural changes based on the alignment model.

According to I-TASSER, amino acid residues at positions 259, 261, 262, 263, 264, 266, 279, 281, 356, 357, 359, 362, 413 and 416 of ROP18 protein sequence were predicted ATP-binding sites. Analysis of residue-by-residue mapping data indicates a high degree of similarity between predicted wild-type ROP18 and known X-ray structure at the active ATP-binding site. However, misalignment of predicted D409A with respect to either predicted wild-type ROP18 or existing x-ray structure at amino acid residues V266 and D362 implies a significant disruption at ATP-binding sites.

Therefore, D409A mutation is hypothesized to cause conformational changes in ROP18 not only on a local level but also on a global scale.

**Table 1.**
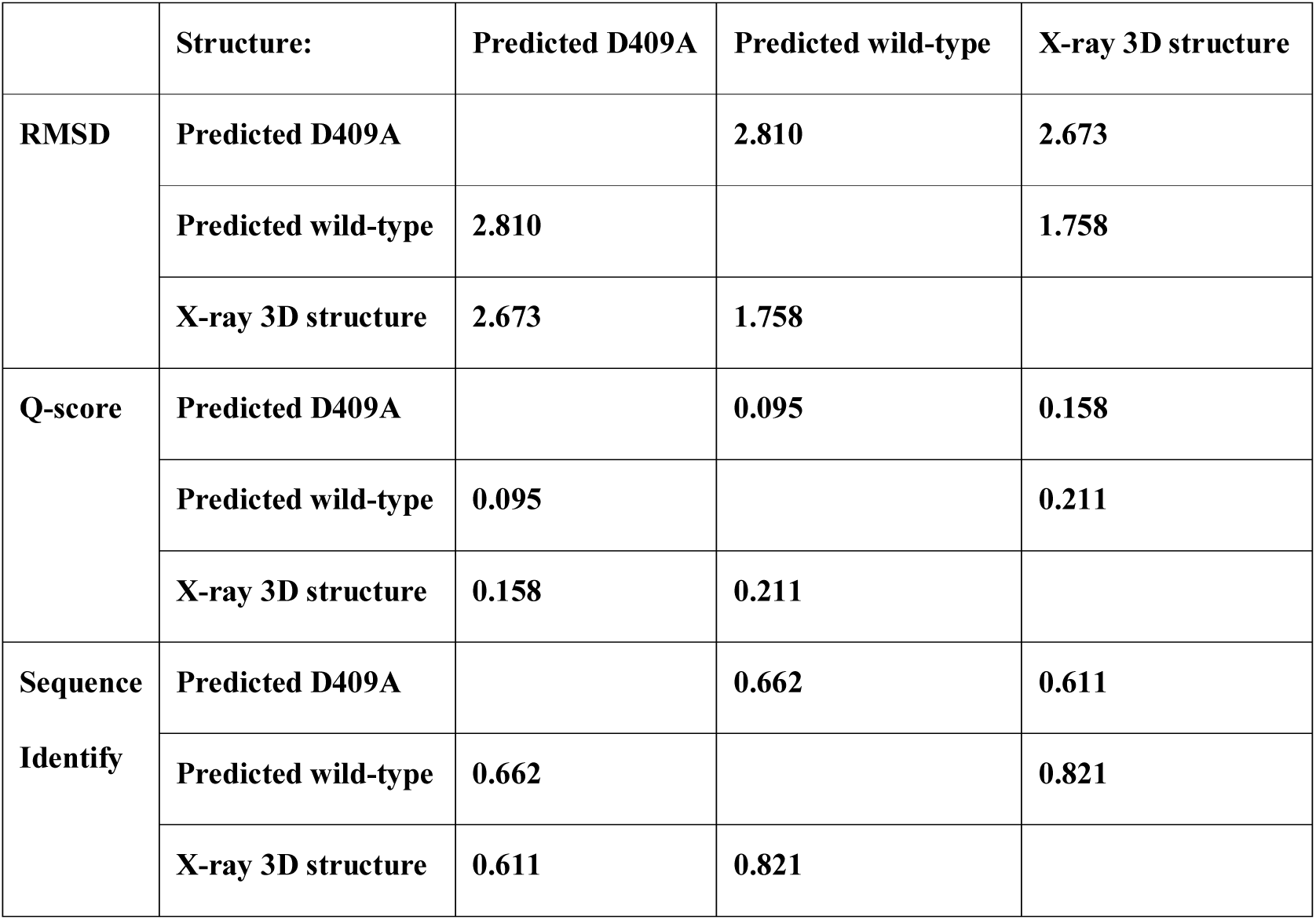
Multiple alignments among predicted wild-type ROP18, predicted ROP18 kinase dead (D409A mutant), and existing ROP18 x-ray 3D structures.

Structural alignments shown were conducted using I-TASSER. RMSD (Root mean square deviation) of aligned structures indicates their divergence from one another. The Q-score takes into account the number of residues in corresponding secondary structure elements (SSEs) and their positions in space. High Q-scores are obtained for structures where a large number of residues in equivalent structural elements superimpose well in three-dimensional space.

### Abrogation of ROP18 kinase activity by point mutation

To characterize the role of the putative catalytic aspartate in the ROP18 protein kinase, site-directed mutagenesis was employed to mutate the amino acid residue to alanine. The necessity of the putative catalytic aspartate residue for kinase activity was explored by comparing the activity of the wild-type ROP18 to that of the D409A mutant. D409A mutation had similar expression level in bacteria to the wild-type ROP18 (Fig 1A). Phosphorylation assays were carried out in the presence of ATP, and reaction mixtures were analyzed by the ADP-Glo™ Kinase Assay Kit. In addition, Western blots using anti-pThr antibody were used to examine the pThr status of the proteins. ROP18 mutants showed a significant reduction in kinase activity, suggesting an inability to be autophosphorylated at the key Thr residues (Fig 1B).

**Fig. 1.**
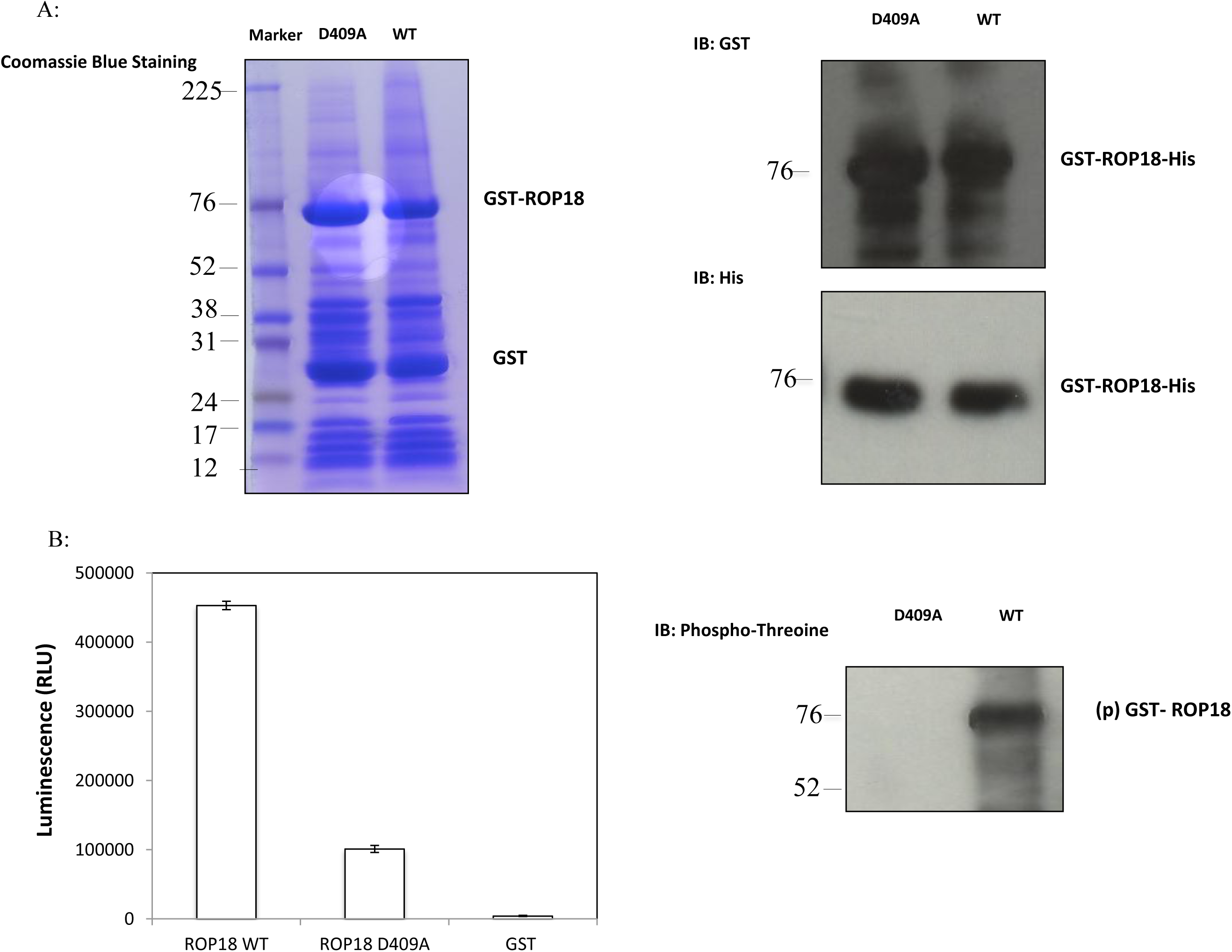
Catalytic aspartate residue is important for ROP18 kinase activity. (A) Expression level of GST-ROP18-His WT and D409A mutant protein in *E.coli*. The expressed proteins were detected by staining with Coomassie Brilliant Blue and analyzed by Western blot using anti-GST and anti-His antibodies. (B) *In vitro* kinase assays were performed using wild-type ROP18 or D409A mutants obtained by recombinant expression in *E. coli* and purification by GST pull-down. Kinase activities were measured by ADP-Glo™ Kinase Assay Kit and also analyzed by Western blot using antibody against pThr.

### Alternation of degradation of ATF6β with ROP18 point mutation

It has been reported that ATF6β is a cellular host target of the *T. gondii* virulence factor ROP18. ATF6β protein is consistently reduced in cells infected with parasites harboring wild-type ROP18 but not with those lacking ROP18 (Δrop18 strain) [21].

To confirm these findings in ROP18-transfected mammalian cells, wild-type ROP18 or D409A mammalian expression vector was constructed. YFP-ATF6β fused to CFP, with a self-cleavage signal [T2A] in between to produce the same level of both proteins, was employed in an ROP18 co-transfection experiment. Protein level of ATF6β was quantified and normalized against ratio of YFP to CFP. There was a moderate reduction of ATF6β protein levels in cells co-transfected with ROP18 kinase dead with respect to the vector control. Reduction in ATF6β protein levels was more significant when cells were co-transfected with wild-type ROP18. Taken together, these data suggest that kinase activity is involved in ROP18-mediated degradation (Fig 2A). In parallel, lysates of 293T cells co-transfected with ATF6β-HA and wild-type ROP18 or ROP18 kinase dead were analyzed by immunoblotting. ROP18-dependent decrease in ATF6β protein level was observed when overexpressing wild-type ROP18 but such phenotype was not present in D409A mutant. Expression of β-Tubulin was consistent across all samples, indicating that general cellular homeostasis was unaffected by ROP18 transfection (Fig 2B).

**Fig. 2.**
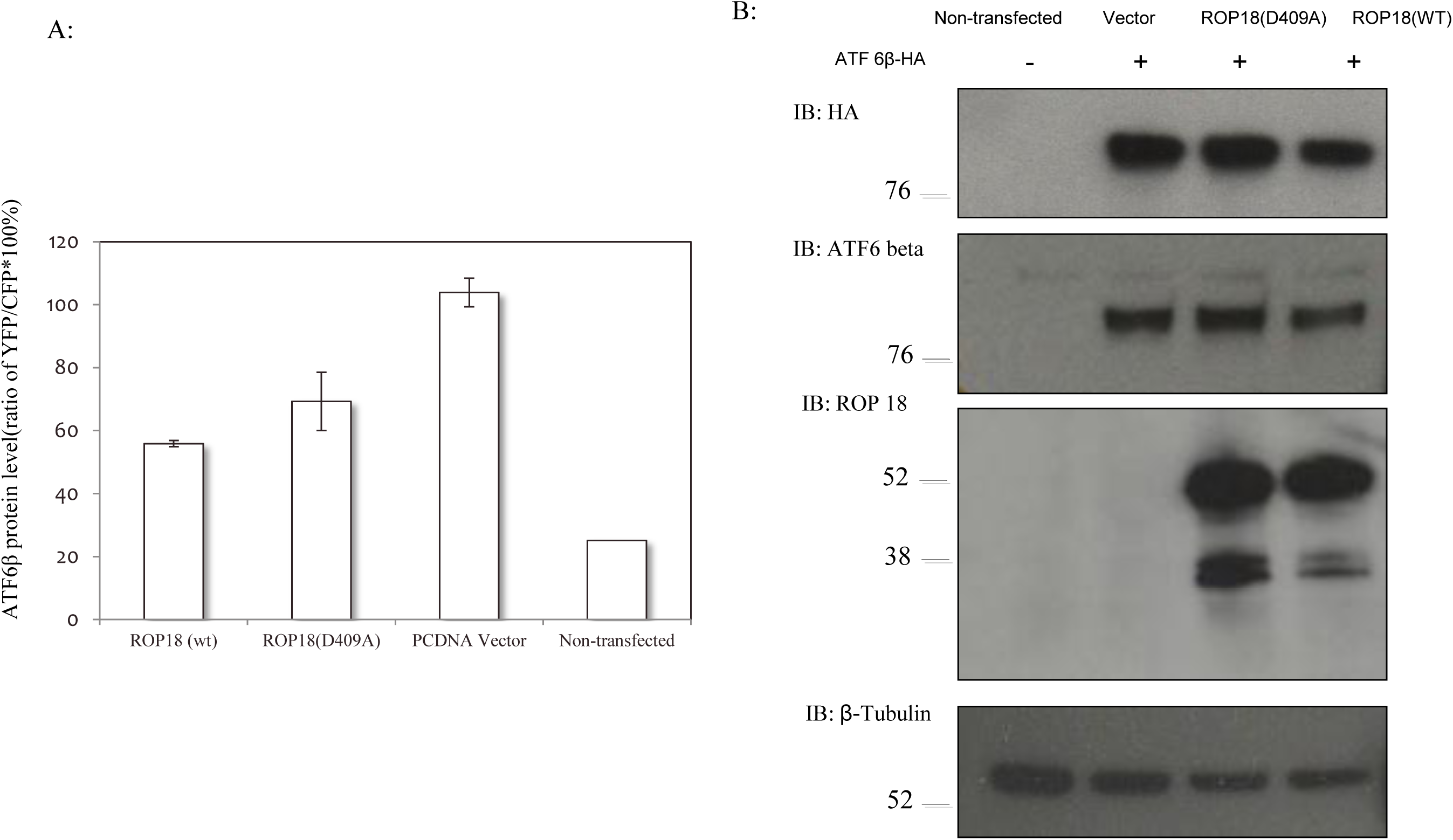
Degradation of ATF6β is associated with and dependent on ROP18 and its kinase activity. (A) Fluorescence readings of 293T cells co-transfected with YFP-ATF6β-CFP and ROP 18. ATF 6β protein expression levels after co-transfecting with either wild-type ROP18 or ROP18 kinase dead. Fluorescence intensity levels of YFP were measured by a fluorescence microplate reader at an excitation wavelength of 500/10 nm and an emission wavelength of 530/10 nm, CFP was measured at an excitation wavelength of 430/10 nm and an emission wavelength of 480/10 nm. Expression levels of ATF 6β were normalized against ratio of YFP to CFP in co-transfected group after 48 hours of transfection. Error bars represent means ± variation range of triplicates. (B) Western blot showing protein levels of co-transfection with ATF6β and either wild-type ROP18 or kinase dead. Cell lysates of co-transfected 293T cells with ATF6β-HA and either wild-type ROP18 or D409A mutant were collocated at 48h post-transfection and analyzed by immunoblotting against HA, ATF6β, ROP18, and β-Tubulin. β-Tubulin was used as a loading control.

To investigate the mechanisms of ROP18-mediated degradation, 293T cells with overexpressed wild-type ROP18 and ATF6β-YFP were treated with MG132, which is a cell-permeable proteasome inhibitor that has been reported to abrogate ubiquitin-mediated proteasomal degradation. Untreated cells were used as a control. Compared with the empty vector control and kinase dead, degradation could be rescued in the presence of MG132, demonstrating that downregulation of ATF6β is proteasome-dependent (Fig 3A). This result was further confirmed by immunofluorescence, where there was a significant increase in ATF6β-YFP signal intensity within ROP18-transfected cells after MG132 treatment in comparison to transfected cells with dimethyl sulfoxide (DMSO) control (Fig 3B).

**Fig. 3.**
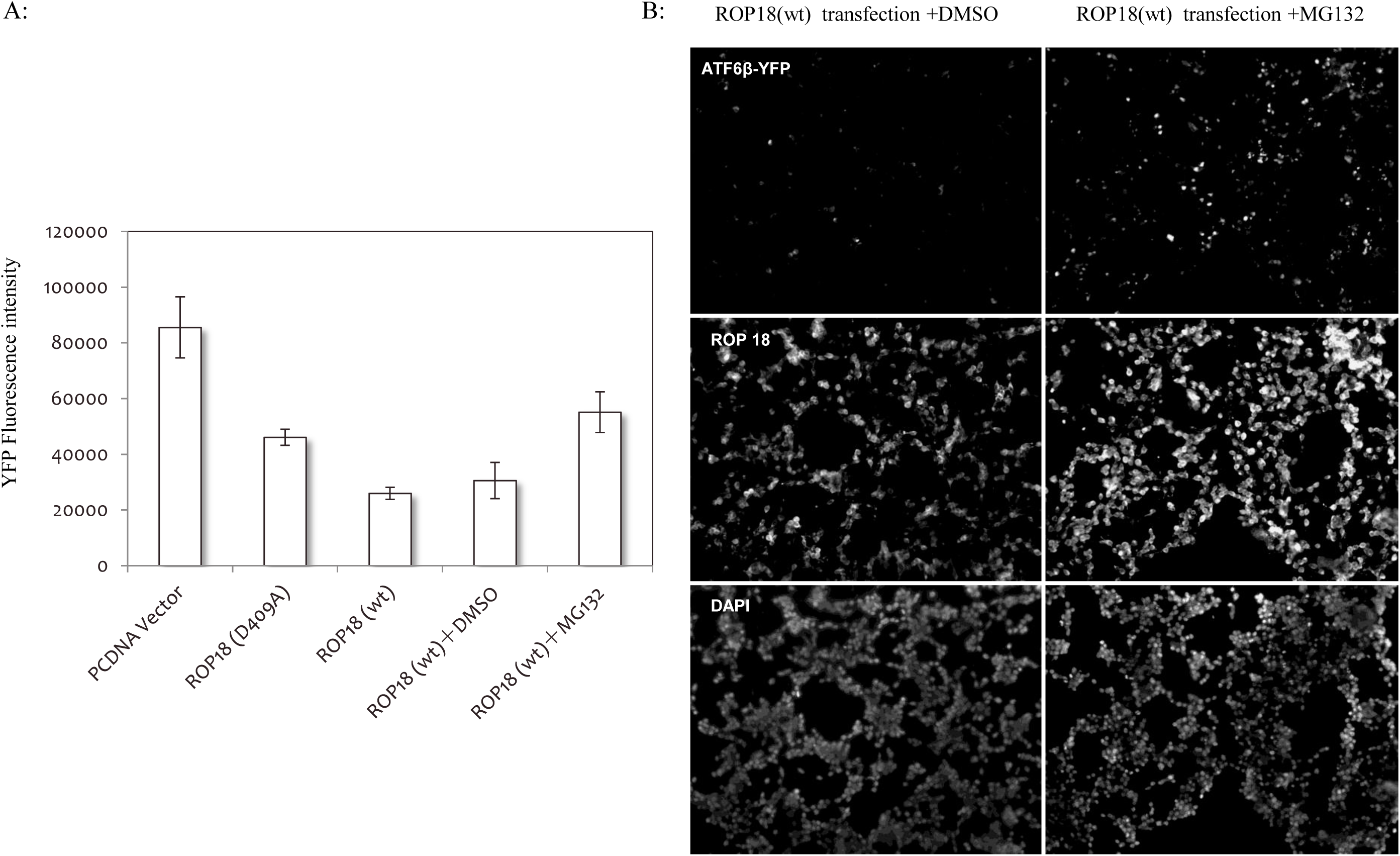
ROP18-mediated degradation of ATF6β is proteasome-dependent. (A) Proteasome inhibitor MG132 rescued ATF6β protein degradation. ATF6β-YFP protein was co-transfected with wild-type ROP18 or ROP18 kinase dead and treated with or without MG132. Fluorescence intensities of YFP were measured by a fluorescence microplate reader at an excitation wavelength of 500/10 nm and an emission wavelength of 530/10 nm. Error bars represent means ± the variation range of triplicates. This blot represents triplicates of two repeats. (B) Rescuing degradation of ATF6β was substantiated by immunofluorescence. ATF6β-YFP protein was co-transfected with wild-type ROP18 into 293T cells with or without 10μM of MG132 treatment. Cells were then fixed, permeabilized, and stained first with rabbit anti-ROP pAb followed by Alexa Fluor^®^ 568 goat anti-rabbit secondary antibody then DAPI. Images were captured with a Leica fluorescence microscope at 10x objectives.

These data are consistent with the previous study, confirming that kinase activity is involved in the ROP18-mediated degradation, and the degradation is in a proteasome-dependent fashion.

### Co-immunoprecipitation assay

To further confirm the physical interaction between ROP18 and ATF6β in transfected 293T cells, cell lysates of co-transfected cells were analyzed by co-immunoprecipitation. Cell lysates of 293T cells transiently transfected with a plasmid encoding HA-tagged ATF6β protein and co-transfected with either wild-type ROP18 or ROP18 kinase dead were prepared. Expressions of recombinant ATF6β-HA and both ROP18 proteins were detected by Western blot using antibodies against ATF6β or ROP18, respectively. Cell lysates were then immunoprecipitated with anti-HA mAb and subsequently immunoblotted with anti-ROP18 pAb to detect the recombinant ROP18. Since wild-type ROP18 mediated degradation of ATF6β, as shown in Fig. 4, ATF6β was reasonably only detected in the sample co-transfected with ROP18 kinase dead (lane 7 of Fig 4 upper panel).

**Fig. 4.**
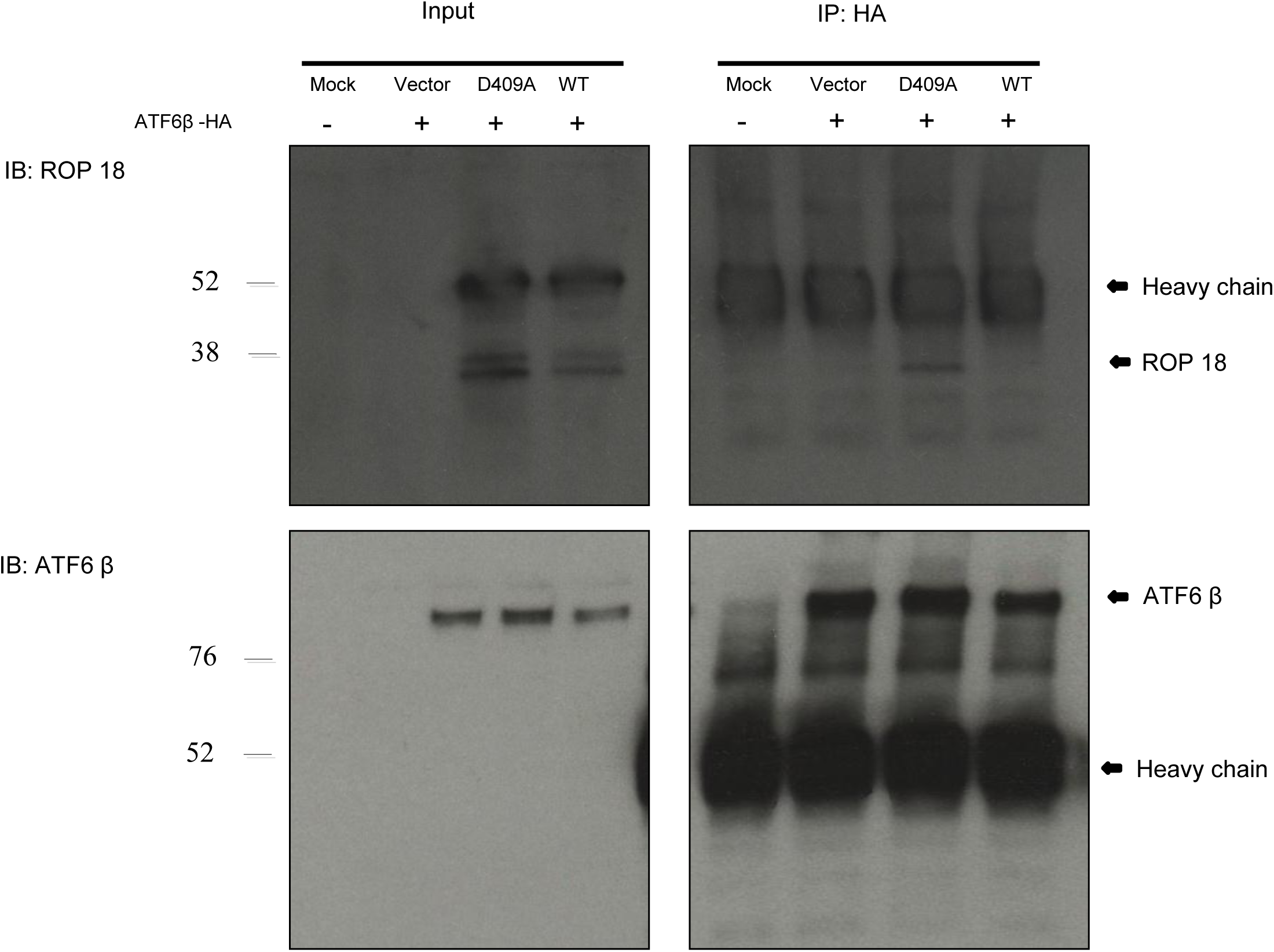
ROP18 and ATF6β physically interact as demonstrated by immunoprecipitation. Lysates of 293T cells transiently co-transfected with 1μg of wild-type ROP18 or ROP18 kinase dead and 1μg of HA-tagged ATF6β expression vectors were immunoprecipitated with antibody against HA and detected by immunoblotting (IB) with antibody against ROP18 or ATF6β. Untransfected and empty vector were used as negative controls.

ATF6β is a member of the basic-leucine zipper (bZIP) DNA-binding protein family, containing an ER-transmembrane domain with the N-terminus facing the cytoplasm [22]. Upon ER stress or unfolded protein response, ATF6β translocates from ER to the Golgi, where its cytosolic domain is released from membrane, after which it translocates once more to the nucleus to activate ER stress-specific genes [23]. The luminal domain of ATF6β is required for sensing ER stress and permitting translocation of ATF6β to the Golgi [26]. Collectively, these results indicated that *T. gondii* ROP18 protein physically binds to recombinant ATF6β in transfected human embryonic kidney cells.

### Subcellular colocalization of ATF6β and ROP18

Colocalization analysis was performed in order to identify where the ROP18-ATF6β interaction occurs in transfected cells.

ATF6 is reported as an endoplasmic reticulum (ER) transmembrane-associated transcription factor [27]. In response to ER stress, ATF6 translocates from the ER to the Golgi, where it is cleaved to be activated [26, 27].

Previous study suggests that overexpression of ATF6 might result in levels of ER chaperones being insufficient for proper folding of exogenous proteins expressed at high levels. As a result, portions of endogenous ATF6 and exogenous ATF6 are subjected to proteolytic processing constitutively [23].

To assess whether overexpressed ATF6β is translocated into Golgi apparatus, 293T cells co-transfected with pcDNA vector control or ATF6β-YFP for 48 hours were fixed for immunostaining with anti-giantin antibody, which is a known Golgi marker. ATF6β-YFP protein co-localized with giantin, thereby suggesting that ATF6β associates with the Golgi (Fig 5, upper panel).

**Fig. 5.**
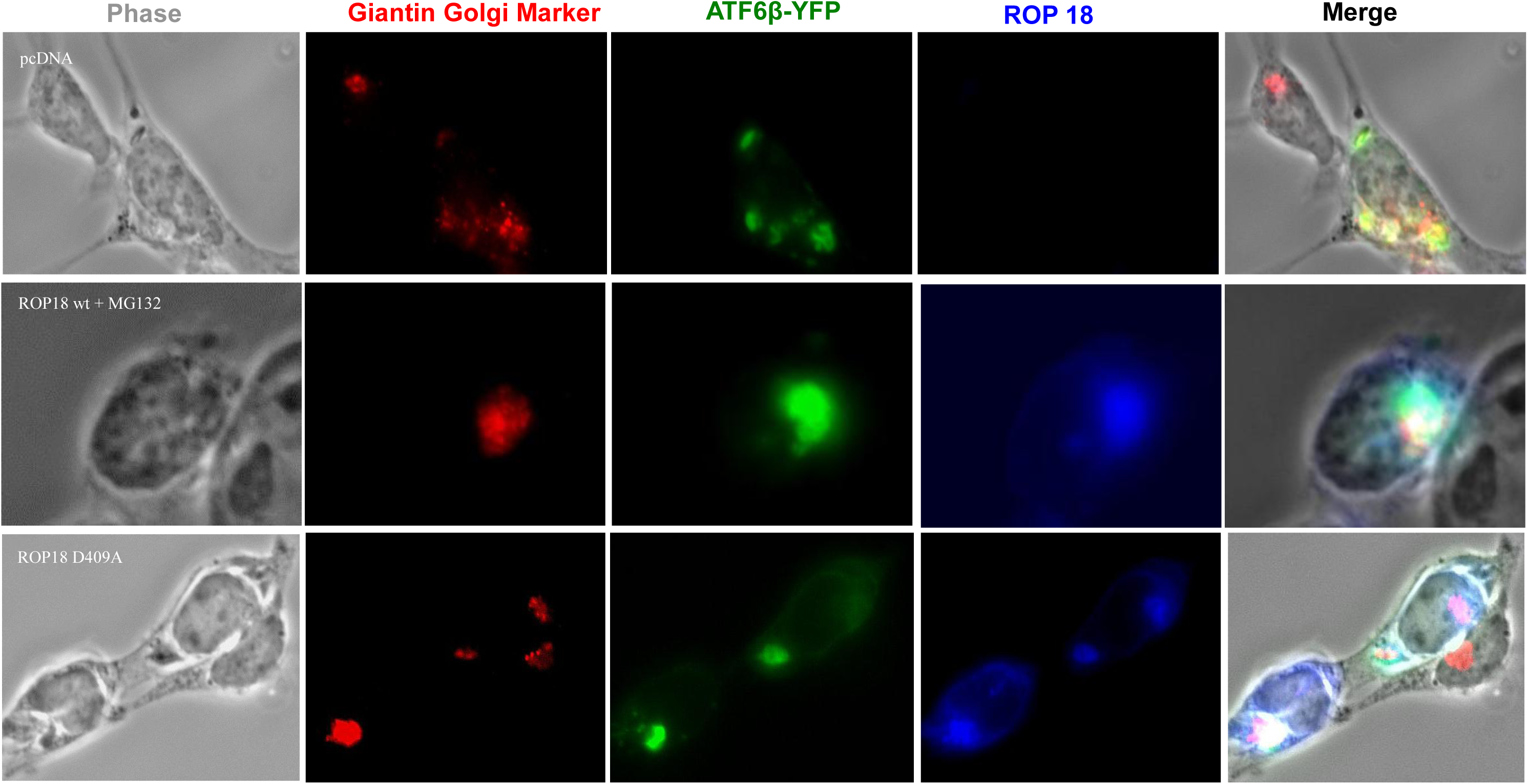
Subcellular colocalization of ATF6β and ROP18 in Golgi compartment within transfected cells. 293T cells co-transfected with wild-type ROP18 or ROP18 kinase dead and ATF6β, with or without MG132 treatment, were analyzed by immunofluorescence assay. Giantin (Red), ATF6β-YFP (Green), ROP18 (Blue) were stained using antibodies specific to giantin (AlexaFluor 568) and ROP18 (Dylight 350) respectively. Images were captured with a Zeiss fluorescence microscope using 63x objectives.

Furthermore, to determine if ROP18 and ATF6β co-localize in the Golgi apparatus, cells co-transfected with ROP18, either wild-type or kinase dead, and ATF6β were analyzed in the same manner. It was established in previous experiments that wild-type ROP18 could mediate ATF6β degradation, and ATF6β could be rescued with MG132 treatment. Cells co-transfected with wild-type ROP18 and ATF6β and treated with MG132, along with cells untreated with MG132 and co-transfected with ROP18 kinase dead and ATF6β, were included in this immunofluorescence analysis. In both conditions, ROP18 was mainly distributed in the Golgi apparatus, where it colocalised with ATF6β-YFP (Fig 5, lower panel).

Altogether these results suggest that *T. gondii* ROP18 protein physically interacts with ATF6β in transfected human cells and that such interaction occurs in the Golgi apparatus.

### Alternation of subcellular localization of ROP18 protein with single amino acid mutation

Intriguingly, while most ROP18 protein was present in the Golgi, some was evident on the plasma membrane (Fig 5, lower panel). To further investigate subcellular localization of ROP18 protein, immunofluorescence staining was performed on cells transfected with wild-type ROP18 and ROP18 (D409A), with or without permeabilization, 24 and 48 hours post-transfection. At 24h post-transfection, both wild-type ROP18 and ROP18 (D409A) were distributed both along cell membrane and in an intracellular compartment close to the perinuclear region – the Golgi compartment. Nevertheless, at 48 h post transfection of ROP18 kinase dead, ROP18 was not only found inside Golgi compartments and on cell surface but also observable in small cytosolic patches, which could be intracellular trafficking vesicles (Fig 6). An inability of those cytosolic patches to colocalize with mitochondrial, lysosomal-associated membrane protein 1 (LAMP1) and Rab11 (a recycling endosomes marker) suggests that said cytosolic patches might be secretory vesicles (S1 Fig).

**Fig. 6.**
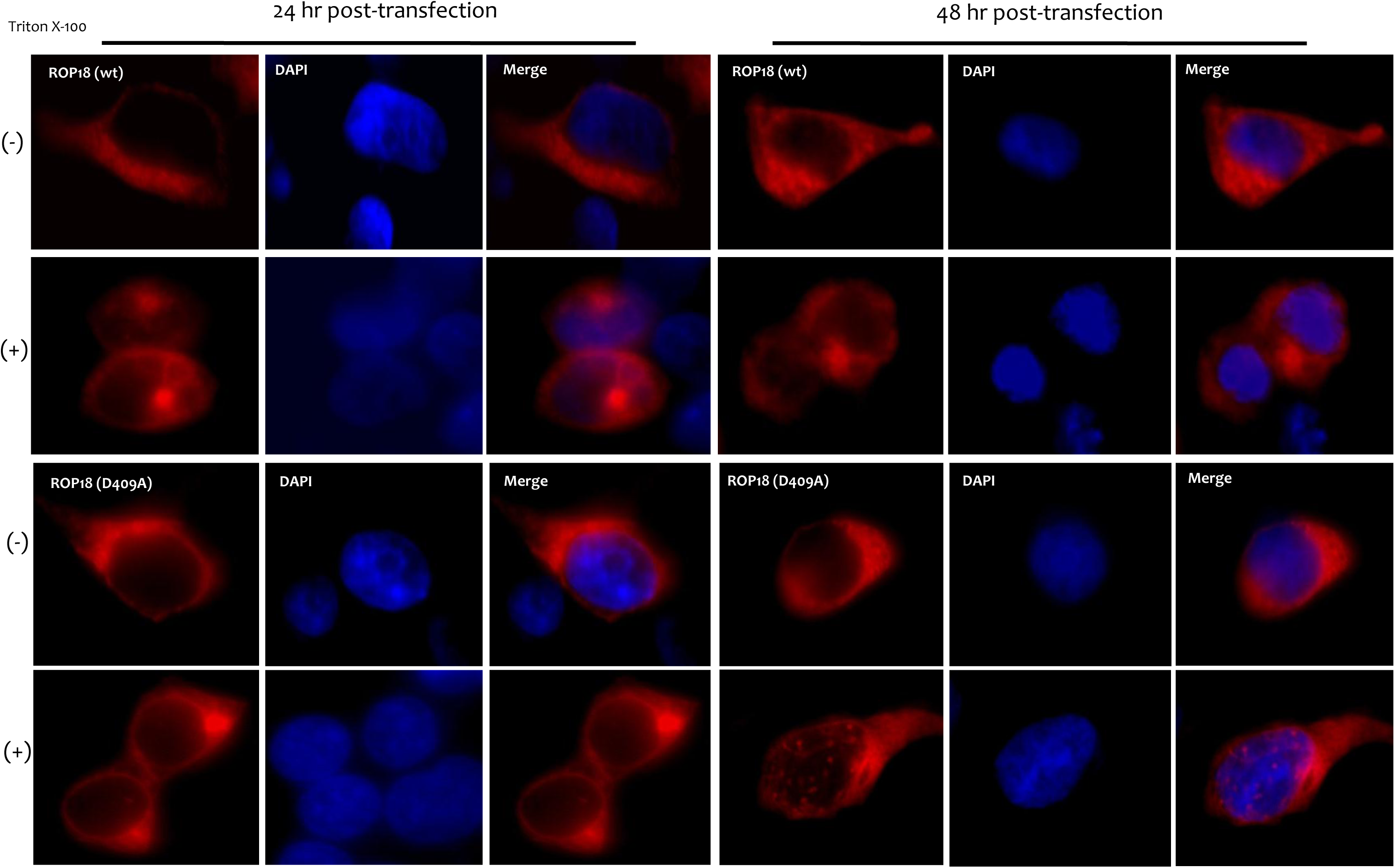
Subcellular localization of ROP18 in transfectd 293T cells. Cells were transfected with wild-type ROP18 or ROP18 kinase dead plasmid for 24hr and 48 hr. Cells were fixed, permeabilized or not with Triton X-100 and stained with rabbit anti-ROP pAb and followed by Alexa Fluor^®^ 568 goat anti-rabbit secondary antibody. Images were captured with a Zeiss fluorescence microscope using 63x objectives.

To ascertain whether those cytosolic patches are secretion vesicles, wild-type ROP18-and ROP18 kinase dead-transfected cultures’ supernatants were collected, concentrated by Amicon concentrators, then analyzed by SDS-PAGE followed by Western blot with antibody against β-Tubulin or ROP18. Cell pellets were collected as control to monitor transfection. Interestingly, ROP18 was present in supernatant of ROP18 kinase dead-transfected culture but not in supernatant of wild-type ROP18-transfected cells. This indicated that ROP18 D409A was secreted. Failure to detect β-Tubulin protein in all supernatants suggested no contamination of intracellular protein (Fig 7).

**Fig. 7.**
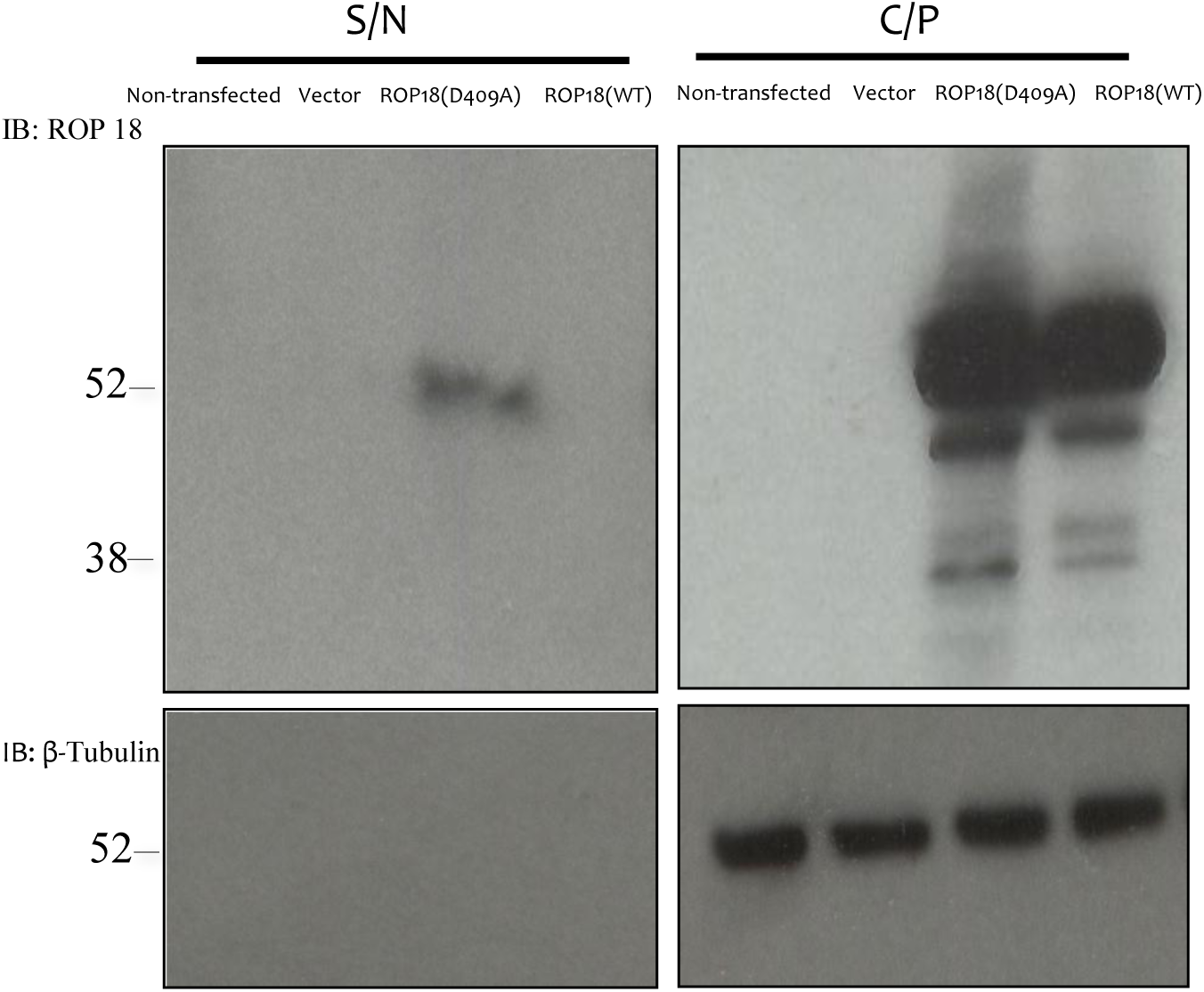
ROP18 kinase dead is released into culture supernatant. Supernatant from cell culture is abbreviated as S/N; C/P is cell pellets. Detection of ROP18 protein in cell culture medium was performed with antibody against ROP18. HEK 293T cells were transfected with empty vector, ROP18 kinase dead or wild-type ROP18 plasmid. Cell culture medium and cell pellets were collected. Culture medium was centrifuged at 13000xg for 5 min to remove the cell debris and concentrated by a 30 kDa-cutoff Amicon concentrator and analyzed by Western blot.

## Discussion

Structural homologies among predicted structures of wild-type ROP18 and ROP18 kinase dead, as well as the known x-ray structure of ROP18, were analyzed by PDBeFold, which is an interactive service for comparing three-dimensional protein structures. Misalignment of protein structures suggests little structural similarity between wild-type ROP18 and ROP18 kinase dead. Furthermore, this point mutation might cause conformational changes not only on a local level, particularly at the proton transport catalytic aspartic acid, but also on a global level in the ROP18 protein.

Site-directed mutagenesis was employed to mutate the critical active site aspartate to alanine to create the D409A mutation. As expected, the single amino acid mutation did not significantly change the protein expression level. On the other hand, the D409A mutant displayed a significant reduced kinase activity *in vitro* and it showed an inability to be autophosphorylated at the key threonine residues.

To better characterize wild-type ROP18 and ROP18 kinase dead, ATF6β protein level was consistently reduced in ROP18 co-transfected cells. In addition, reduction in ATF6β protein levels was more significant in cells co-transfected with wild-type ROP18 compared to those co-transfected with ROP18 kinase dead, suggesting that kinase activity is involved in ROP18-mediated degradation. Proteasome inhibitor MG132 was able to rescue ROP18-associated degradation, suggesting that downregulation of ATF6β process is proteasome-dependent.

These data confirm that kinase activity is indispensible for ROP18-mediated degradation, and the point mutation at key catalytic residue could abolish ROP18’s function.

Cells have developed signaling pathways to monitor folding environment of the ER and adjust the ER folding capacity accordingly. When unfolded or misfolded proteins accumulate in the ER, signals are transmitted from the ER to the nucleus and cytoplasm. These pathways are termed the ER stress or unfolded protein response. ATF6β, a primary ER stress sensor, initiates several cellular responses and signaling pathways to restore ER homeostasis.

*T. gondii* parasitophorous vacuole membrane (PVM) was reported to form tight associations with host mitochondria and the ER. Therefore, it is plausible that ER-localized ATF6β in vicinity of PVMs might gain access to ROP18. Indeed, ATF6β-YFP and CFP-tagged ROP18 kinase dead were colocalized with ER-specific RFP in transfected 293T cells, suggesting that ROP18 is in close proximity to proteins localized in the ER. HFFs infected with parasites were analyzed by electron microscopy, pointing out that PVMs directly fuse with the host ER [21].

Earlier research suggests that *T. gondii* infection could trigger unfolded protein response [28]. Previous study also demonstrates that overexpressing HA-ATF6β results in activation of unfolded protein response in 293T cells; however, co-expression of ROP18 downregulates ATF6β-dependent activation in a dose-dependent manner [21].

In response of ER stress, ATF6 translocates from the ER to the Golgi, where it is cleaved and processed to its active form [26, 27]. ATF6 has been shown to be processed by Site-1 protease (S1P) and Site-2 protease (S2P), both of which require RxxL and asparagine/proline motifs [29]. Coincidentally, ROP18 sequence has RMGL and NREP motifs, both of which might be recognized and processed by S1P and S2P at Golgi. ROP18 does undergo cleavage, and its post-cleavage form was detected by the polyclonal antibody against ROP18, as shown in Fig 2B. The post-cleavage ROP18 interacted with ATF6β protein in the co-transfected cells. Taken together, these data suggest that ROP18 might migrate to the Golgi and also interact with ATF6β.

Colocalization of ROP18 kinase dead and ATF6β confirmed that the interaction might also occur in the Golgi apparatus. Overexpressed ROP18 could translocate into Golgi, where ATF6β is known to be cleaved and processed, suggesting that this transient interaction starts from ER to the Golgi. Since the Golgi serves as a major protein-sorting hub for secretory cargos, it receives *de novo* synthesized proteins from the ER destined for secretion from the cell or for residence in endocytic branches of the secretory pathway or along the plasma membrane [30]. In order to determine whether overexpressed ROP18 could migrate to cell membrane from the Golgi, immunofluorecence staining on ROP18 transfected cells, with or without permeabilization, was performed. Association of ROP18 with cell membrane was observed, suggesting a membrane-targeting strategy for ROP18.

Being so instrumental to *T. gondii* pathogenesis, then ascent parasitophorous vacuole membrane is believed to derive from host cell plasma membrane [31]. This fact is consistent with the finding that ROP18 localizes to early- and late-stage parasitophorous vacuole membranes [32]. In addition to cell membrane localization, some ROP18 kinase dead was found located inside intracellular trafficking vesicles, and further experiment demonstrated that some ROP18 kinase dead was secreted into culture medium.

Whether there is any correlation between eliminating kinase activity and localization or secretory pattern remains to be elucidated. However, it is plausible that altering kinase activity itself or changing protein hydrophobicity and hydrophilicity as a result of mutations engenders protein mislocalization.

Collectively, findings presented herein demonstrate that a single amino acid mutation on the proton transport catalytic aspartic acid induced functional changes associated with ROP18 protein, and these changes are very likely associated with conformational alterations in the secretory protein.

