## Supplementary Materials for "Deactivation and mislocalization of a kinase protein induced by a single amino acid mutation on the proton transport catalytic aspartic acid" (from another version of this preprint: version 1, 2018-03-15)

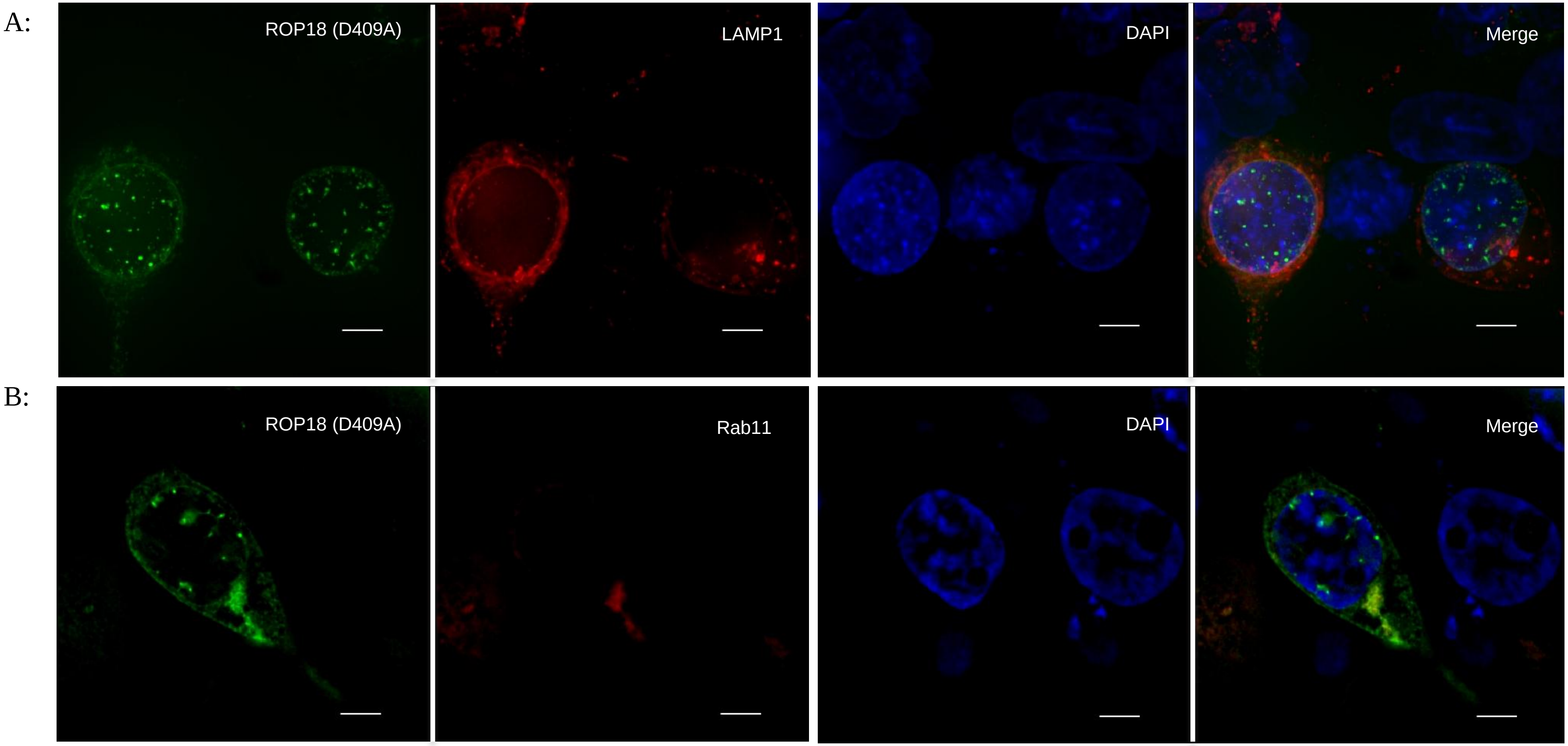


**S1 Fig. Subcellular localization of ROP18 in transfectd 293T cells.** Cytosolic patches did not colocalize with lysosome and recycling endosomes. Cells were transfected with ROP18 D409A and either LAMP1-mcherry (A)or Rab11-mcherry (B) for 48 hours. Cells were fixed, permeabilized and stained either with rabbit anti-ROP18 mAb and followed by Alexa Fluor® 488 goat anti-Rabbit and DAPI staining.
